# Fast and cost-effective age estimation in Bengal tiger and Asiatic lion: applicability of cementum analysis method

**DOI:** 10.1101/2021.09.27.461978

**Authors:** Vipin, Chandra P Sharma, Vinita Sharma, Surendra P Goyal, Sandeep K Gupta

## Abstract

Age estimation methods, through cementum analysis, for wild animals are rarely developed in Southeast Asian Countries. In the present study, we describe the applicability of the cementum analysis technique for developing a fast (±19 hours) and cost-effective age estimation method for Bengal tiger (*Panthera tigris tigris*) and Asiatic lion (*Panthera leo persica*) using incisor tooth. The I_2_ and I_3_ incisor teeth from the right mandible of a tiger and I^2^ and I^3^ from the left maxilla of a lion were used in the study. The longitudinal sections of the tooth were made using a low cost hand grinding technique on sand papers followed by decalcification and staining with hematoxylin. The cementum layers were counted under the microscope at 100X or 200X magnifications. Two cementum layers were observed in each of the I_2_ and I_3_ incisor tooth of tiger and six cementum layers were observed in each of the I^2^ and I^3^ incisor teeth of lion. The permanent incisors in tiger and lion erupt between 12-14 months; hence, we added 1 year to the counted number of cementum layers to estimate the final age of tiger and lion incisors. The absolute age of tiger and lion incisors was estimated to be of 2+1 years and 6+1 years, respectively. The same number of cementum layers in both incisors respective to the tiger and lion were observed. Therefore, we suggest (i) undertake the blind test and (ii) collect incisor teeth from naturally died or killed individuals for strengthening the database on the age of the wild population. This optimized method may be suitable for many carnivore species, applicable in wildlife forensic studies and can be used by researchers with minimum expertise, time, and funds requirements throughout the world.

## Introduction

The age estimation methods in big cats are required for studies related to the species demography for understanding population dynamics (Foresman et al. 2012; Skalski et al. 2005), age class (Creel et al. 2004, Angerbjorn et al. 2004), population monitoring trends (Barthold et al. 2016), negative human-wildlife interactions (Conover 2002, Thirgood & Rabinowitz 2005), and in illegal wildlife trade (Williams et al. 2015). Among the most common methods available for carnivores age estimation are tooth eruption (Slaughter et al. 1974), wearing of a tooth crown (Harris 1978, Stander 1997, Gipson et al. 2000), closure of pulp chamber (Marks and Erickson 1966, Zapata et al. 1997, Binder and Van Valkenburgh 2010), and cementum analysis (Klevezal & Kleinenberg 1967, Matson 1981, White & Belant, 2016).

In tiger, the age estimate analysis has mostly been limited to tooth eruption and wearing (Mazak 1979 & 1981, Miles & Grigson 2003). For lions, the methods described are sizes of body and mane, pigmentation in nose and tooth wear (Schaller 1972, Smuts et al. 1978, Whitman et al. 2004, Whitman and Packer 2007, Ferreira & Funston 2010), closer of the pulp chamber (White & Belant 2016), the ratio of tooth areas (White et al. 2016), tooth eruption (Schneider 1959) and cementum analysis (Spinage 1976, Smuts et al. 1978; White & Belant 2016).

Amongst the various age determination methods available, the cementum analysis method has been recommended for its accuracy (Matson et al. 1993, Klevezal & Kleinenberg 1967, Johnston et al. 1987, Mundy & Fuller 1964, Marks & Erickson 1966, Craighead et al. 1970, Willey 1974, Grue & Jensen 1979, Mbizah et al. 2016, Vipin et al. 2018). However, all the above methods, except Vipin et al. (2018), are costly and time-consuming, limiting their adaptability by researchers working in the labs with limited resources and funds. Moreover, the expertise required in all these techniques needs special training for the users.

Rare studies are available in Southeast Asia for wild animals age estimation through cementum layers analysis (Vipin et al. 2018) and non for a carnivore species. Therefore, in the current study, we present the applicability of the cementum analysis technique to develop a fast and cost-effective age estimation method for tiger (*Panthera tigris tigris*) and Asian lion (*Panthera leo persica*). The technique does not need a costly microtome for tooth sectioning, so the most of the items required are generally available in a common lab. The current study may be applicable to other carnivore species requiring less time, expertise and money.

## Material and Methods

### Sample collection and preparation of longitudinal section of tooth

The two permanent incisor teeth (I_2_, I_3_) out of three were used from the right mandible of a tiger (R-5370) (Fig. 1A, B and C) and two incisors (I^2^, I^3^) from the left maxilla of a lion skull available at Wildlife Forensics and Conservation Genetics Cell, Wildlife Institute of India, Dehradun were used in the study. The teeth from the tiger mandible were extracted by boiling it in water for ten minutes, after which they detach themselves from the mandible. From the lion skull, the teeth were removed with the help of a plier with utmost care that the periodontal membrane remains intact. We used our earlier described protocol of preparing the longitudinal sections of the incisor tooth (Vipin et al. 2018), as shown in Fig. 2.

**Figure 1.**
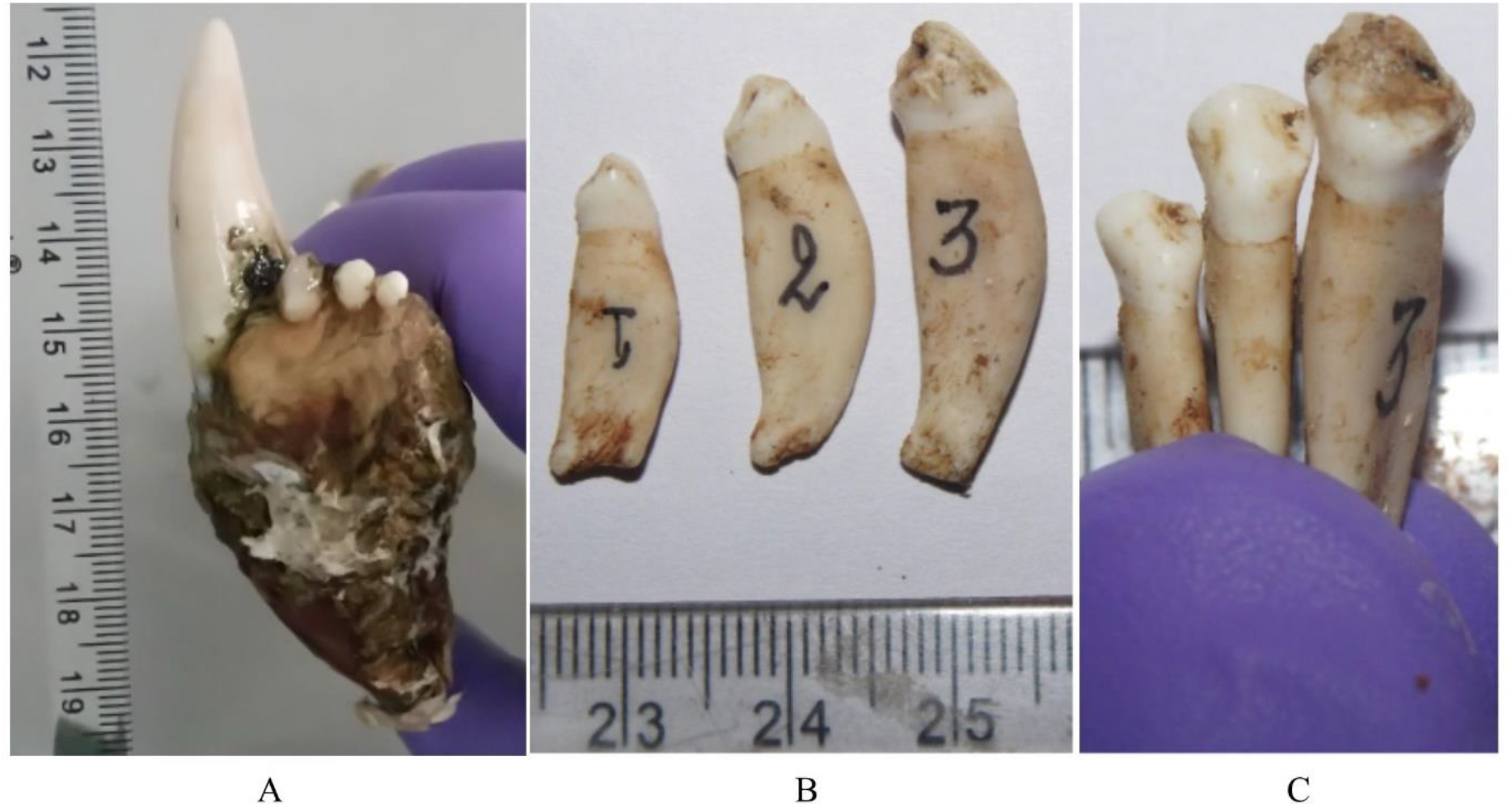
Images showing tiger mandible (A), extracted incisors (I_1_, I_2_, I_3_) in side view (B) and lingual view (C).

**Figure 2.**
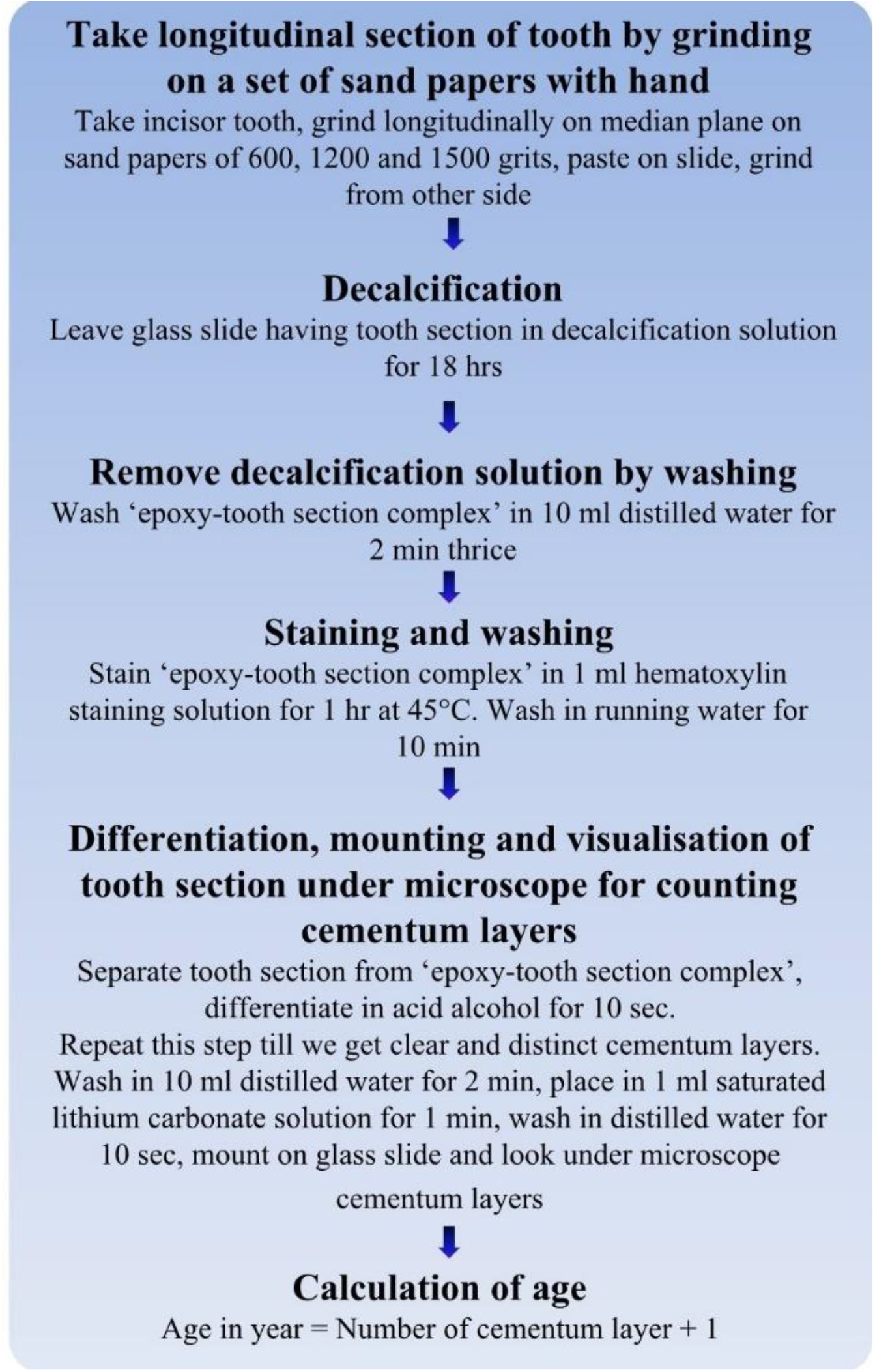
Flow chart of the procedure followed for determination of age through cementum layer count in incisor tooth.

### Calculation of age from cementum annuli

We counted the acellular cementum layer in the root portion of the tooth, which is formed annually and stains dark with hematoxylin (Matson et al. 1993). The permanent incisors in tiger and lion erupt completely between 12-14 months (Smuts et al. 1978; Mazák 1979 and 1981). Therefore, we added a minimum of one year in the final age estimation calculations for both species. The presence of one cementum layer in a permanent incisor tooth of tiger and lion indicates that the animal has lived one year, at least. Therefore, the age of sectioned tooth in years was calculated according to the formula

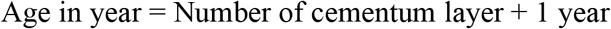

## Results

The teeth sections of the tiger showed two cementum layers for I_2_ (Fig. 3 A, B, and C) and I_3_ (Fig. 4A and B). Therefore, the final age of the tiger was estimated by adding 1 year to the counted number of cementum layers which came out to be 2+1 years. The cementum layers in I^2^ (Fig. 5A and B) and I^3^ (Fig. 6) from the left maxillae of the lion came out to be six. Therefore, the final age of the lion was estimated to be of 6+1 years. The cementum layers in the incisor tooth were photographed wherever these were seen distinct and clear.

**Figure 3.**
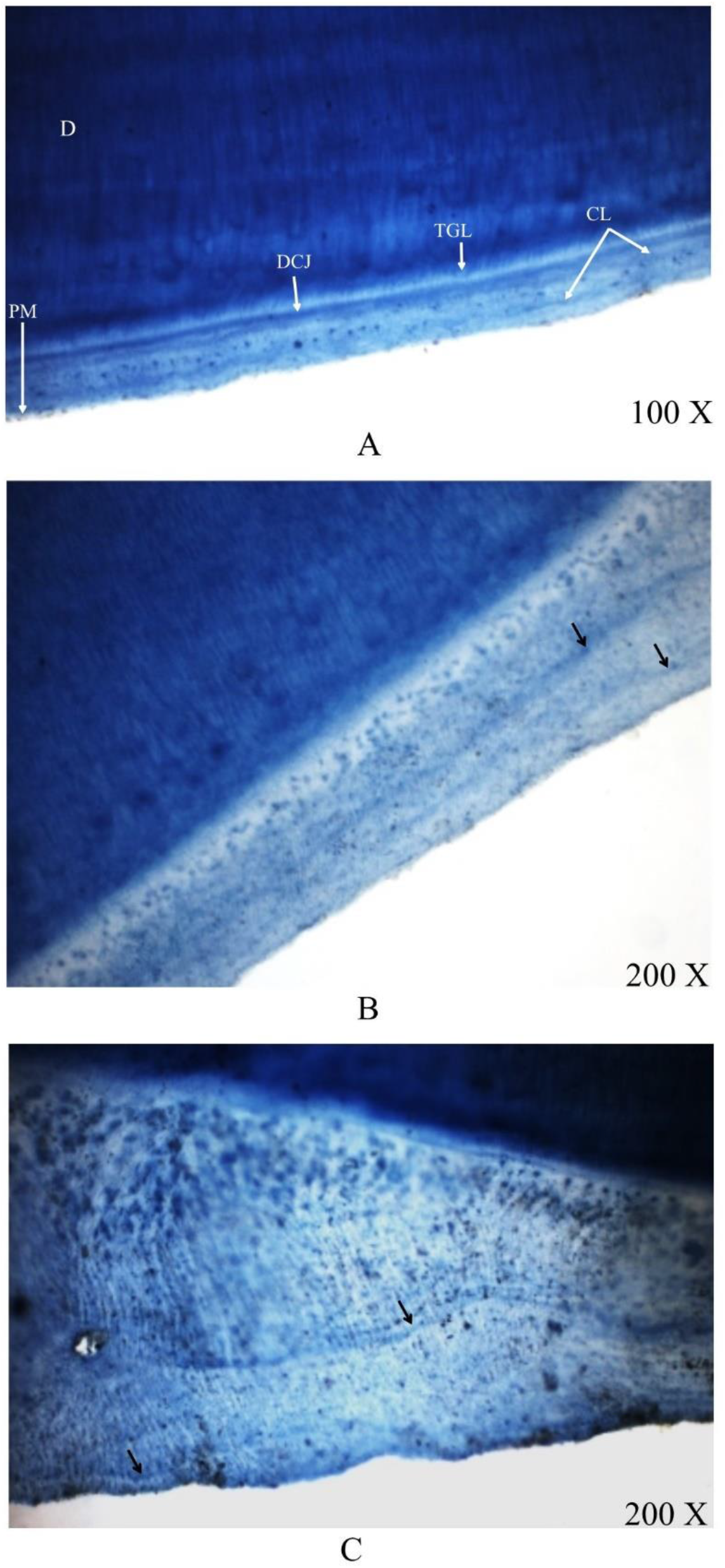
The longitudinal section of I_2_ tooth of tiger showing two cementum layers in three different regions (A, B and C) under 100X and 200X magnifications. D=dentine, TGM=Tome’s Granular Layer, CL=Cementum Layer, DCJ=Dentine Cementum Junction, PM=Periodontal Membrane.

**Figure 4.**
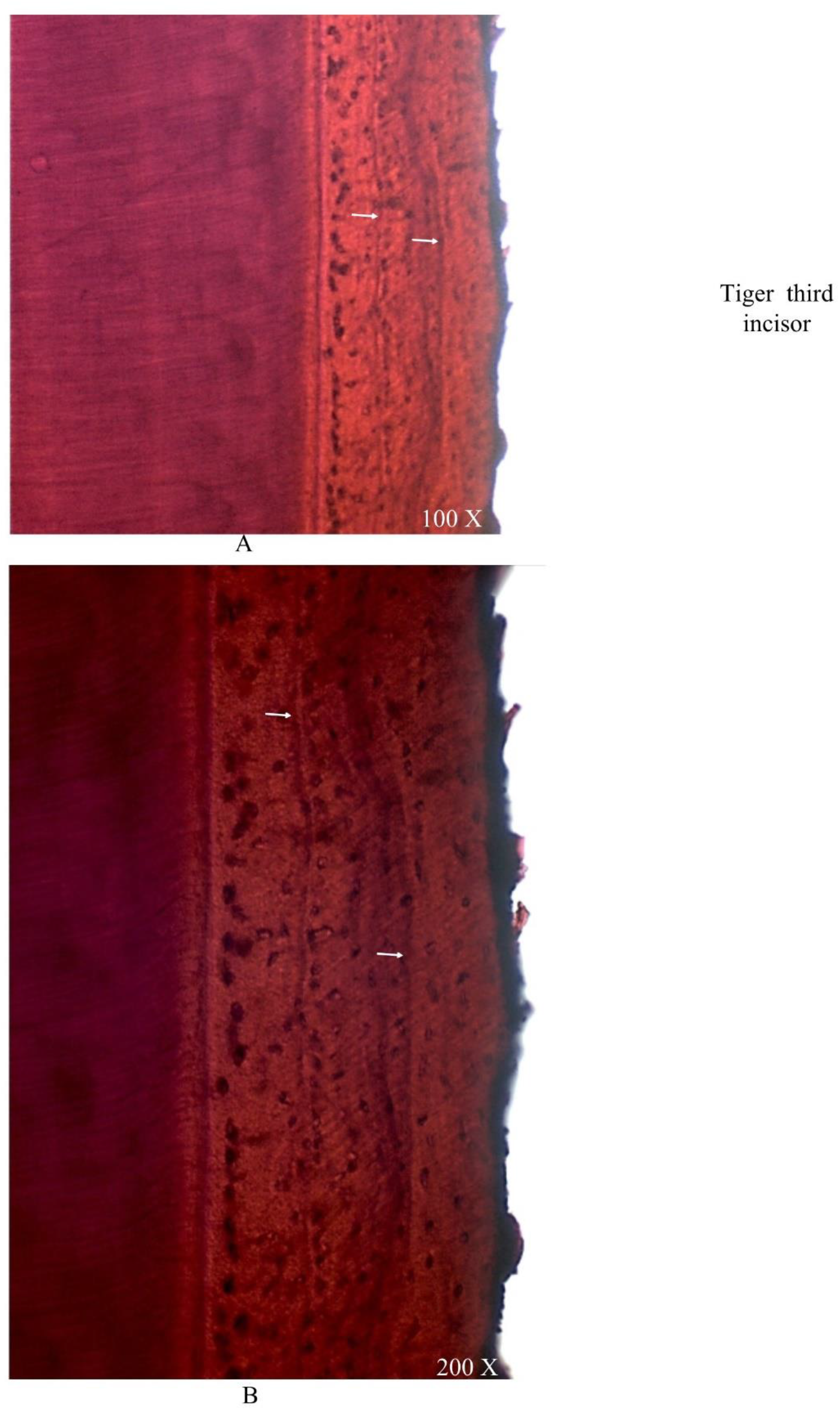
The arrows in the longitudinal section of I_3_ of the tiger showing two cementum layers at 100X (A) and 200X (B) magnifications.

**Figure 5.**
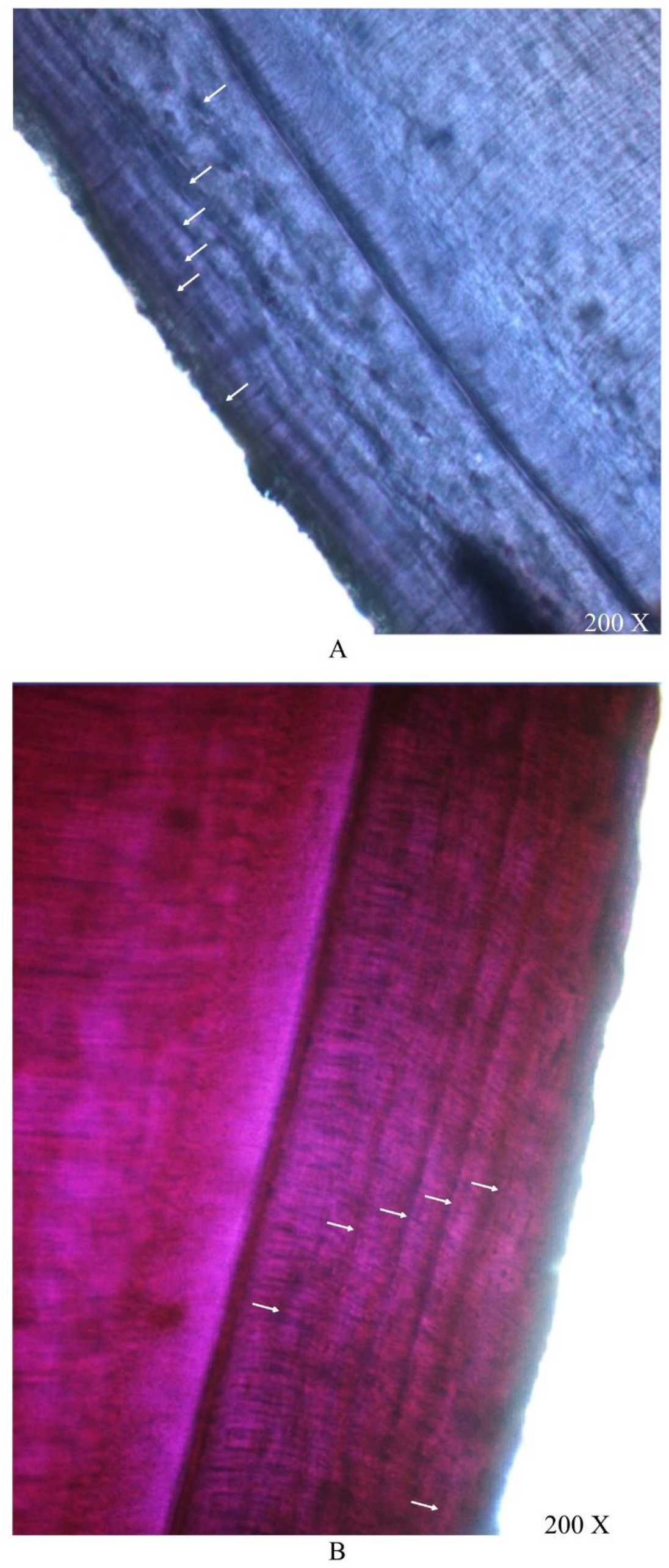
The arrows in the longitudinal section of I^2^ of the lion showing six cementum layers in two different regions (A and B).

**Figure 6.**
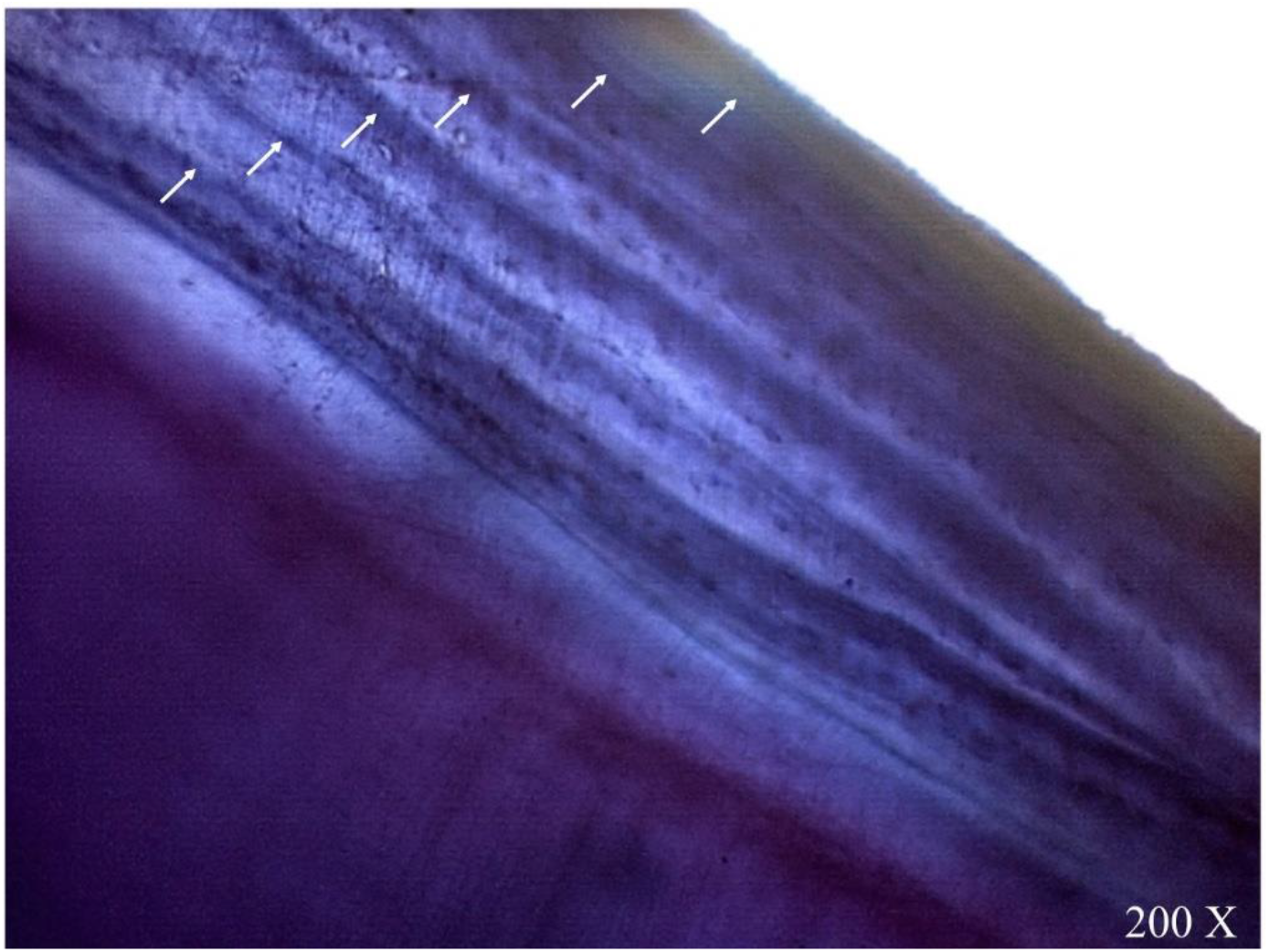
The arrows in the longitudinal section of I^3^ of the lion showing six cementum layers.

## Discussion

The methodology used here for the estimation of tiger and lion ages has been adopted from our previous study related to the *Axis axis* age estimation (Vipin et al. 2018), where the quality, accuracy, low time consumption in comparison to other similar techniques, its cost effectiveness, advantages, disadvantages, precautions, troubleshooting and other, related, important points, in general about the method have already been discussed in detail. In our earlier study, it has been proved that the thickness of the longitudinal section (around 57 μm) was adequate to show all structures in a tooth which are required for the estimation of age, the results were comparable to a microtome procedure and the cementum layers counted for *Axis axis* were as accurate as counted by Matson’s Laboratory (Vipin et al. 2018). However, the species-specific parameters viz. tooth type, diet, nutrition, climatic conditions of their geographic factors do matter a lot while developing age estimation protocol using cementum analysis for any species. Here, we have tried to address all these concerns with respect to the tiger and lion. The incisor teeth are the first to erupt in permanent dentition in tiger and lion (Smuts et al. 1978, Mazak 1981, Currier 1983).

In cats, all permanent incisors except I^3^ erupt before other teeth (Colyer 2003). In tiger, the permanent tooth eruption starts between 8.8-9.5 months and completes between 12-14 months (Mazák 1979 and 1981). In lion, permanent I^1^, I_1_, I^2^, I_2_ erupts completely between 9-11 months and I^3^ and I_3_ start erupting by the end of this period (Smuts et al. 1978). I^3^ and I_3_ completely replace their deciduous counterparts by 12-14 months, while P^2^ starts erupting between this period in the lion (Smuts et al. 1978). Though no published data related to age estimation of tiger using cementum analysis is available, many researchers have used permanent incisors, canine, and second premolar tooth to develop age estimation methods in lions utilizing this technique (Smuts et al. 1978, Cheater 2006, White & Belant 2016). In tiger and lion, the canines are more in demand than other tooth types in the illegal wildlife trade, so they might not be available for determining age. In both species, the permanent incisors number is six times more than P^2^, which is a plus point if some tooth gets damaged during processing for cementum analysis. Therefore, we selected incisors in the current study and based on the availability of their types; the incisors were extracted from the skull. Hence, the criteria of keeping a common incisor type from a particular mandible or maxilla was not possible.

The time taken by different tooth types for their permanent eruption has been reported unequal in other species of carnivores and ungulates (Zapata et al. 1995, Azorit et al. 2004). There is no information available for the first rest line formation in tiger and lion incisor tooth. In P^2^ of African lions, it is established that the first rest is formed in the second year of age, so the authors had to add 1 to the counted number of cementum lines to estimate the final age (White & Belant, 2016). As mentioned above, all permanent incisors in tiger and in African lion are present when the permanent P^2^ erupts in an African lion (Smuts et al. 1978). Therefore, we assumed that the first rest line should be present in the second year of age for both of these species, and hence we added 1 year to the counted number of cementum layers for final age estimation.

Many studies have relied on two or more teeth in the absence of known age animals of a common type, sectioning while confirming the age through cementum layer analysis. It has been shown in a blind duplicate test that first permanent incisors from right and left mandibles of *Axis axis* of unknown age showed the same numbers (n=6) of cementum layers in each tooth (Vipin et al. 2018). White and Belant (2016) have used paired PM^2^ while estimating the age of free-ranging African lions of unknown age through cementum line count as one of the methods in their study. Therefore, before proceeding with this study and in the absence of large sample size and lack of known age teeth for tiger and lion, we hypothesized that at least two permanent teeth, of a common type, of an animal should represent the same number of cementum layers. It is essential to discuss here that White and Balent (2016) have shown that cementum layer count in PM^2^ is unsuitable for aging lions. Their analysis revealed that in most of the PM^2^ pairs (19 out of 31), the cementum line count was differed by 1-2 lines and even went up to 7 lines for other pairs. Smut et al. (1978), while aging African lions through cementum lines in canine tooth, found that this technique gave the results in compliance of the known ages of lions with very high and significant correlation value (*r*=0.973; *P*<0.001). So, to compare the results of White and Balent (2016) with Asian lions, the P^2^ tooth with a large sample size or the data of more incisors or a different tooth type needs to be analyzed for cementum layer count. Otherwise, it would be inappropriate to say anything in this regard about the current study. In ungulates, the accuracy percentage of age estimation through cementum analysis decreases with the increase of age (Hamlin et al. 2000); more research can provide an idea about carnivores. Our concern here is to present a fast and straightforward age estimation method for Bengal tiger and Asiatic lions, which further has a scope of research to address the above points.

The applicability of the cementum analysis technique for tiger estimated it’s age as three years. The two dark cementum layers in both I_2_ and I_3_ indicateed two winter seasons, which means that the animal has lived at least two years. But the acellular cementum region after the second dark line up to the periodontal membrane in both the teeth indicated the period from last winter to July 2016 when the animal died due to an accident. So, according to this, an approximate time of 6 months should also be added to the final estimated age, and it should come around 3 years and 6 months. Since, we did not have a sufficient sample size of known age tigers to establish this fact, we chose to restrict our estimate, here, to the minimum number of years. A robust age estimation methodology cannot be drawn for similar tooth types of other species unless an extensive data set with known age animals is included. We also suggest that the parameters like uneven nutrition, climatic, physiological, and geographic factors should also be analyzed to estimate tiger and lion age using cementum analysis, with additional samples. The information on the demography of a population of a species may be derived from the age estimates of its individuals (Veiberg et al. 2020). Analysis of population dynamics parameters viz. survivability, parenting, reproductive success, size of the body, aging is challenging without the knowledge of a reliable aging method (Berube et al. 1999; Douhard et al. 2016; Hewison and Gaillard 2001; Nussey et al. 2006; Pelletier et al. 2012; Weladji et al. 2002). Therefore, we suggest the collection of incisors from the dead or killed individuals of wild populations.

In previous study, we suggested that this technique has relevance in wildlife forensics when there is a need to know which age class of a particular species is preferred in the illegal wildlife trade (Vipin et al. 2018). So, we tried to estimate the age of an incisor from the mandible of a tiger seized in the illegal wildlife trade and was sent to Wildlife Institute of India, Dehradun, for species confirmation. We found two dark cementum layers for this tooth; hence it’s age was estimated as three years (Fig. 7). Therefore, the developed method may also be applied on large samples size to know which particular age class of tigers are in the illegal wildlife trade. The developed method is fast and cost-effective, it has worldwide application in estimation of age using cementum analysis, especially in laboratories where the resources and funds are limited.

**Figure 7.**
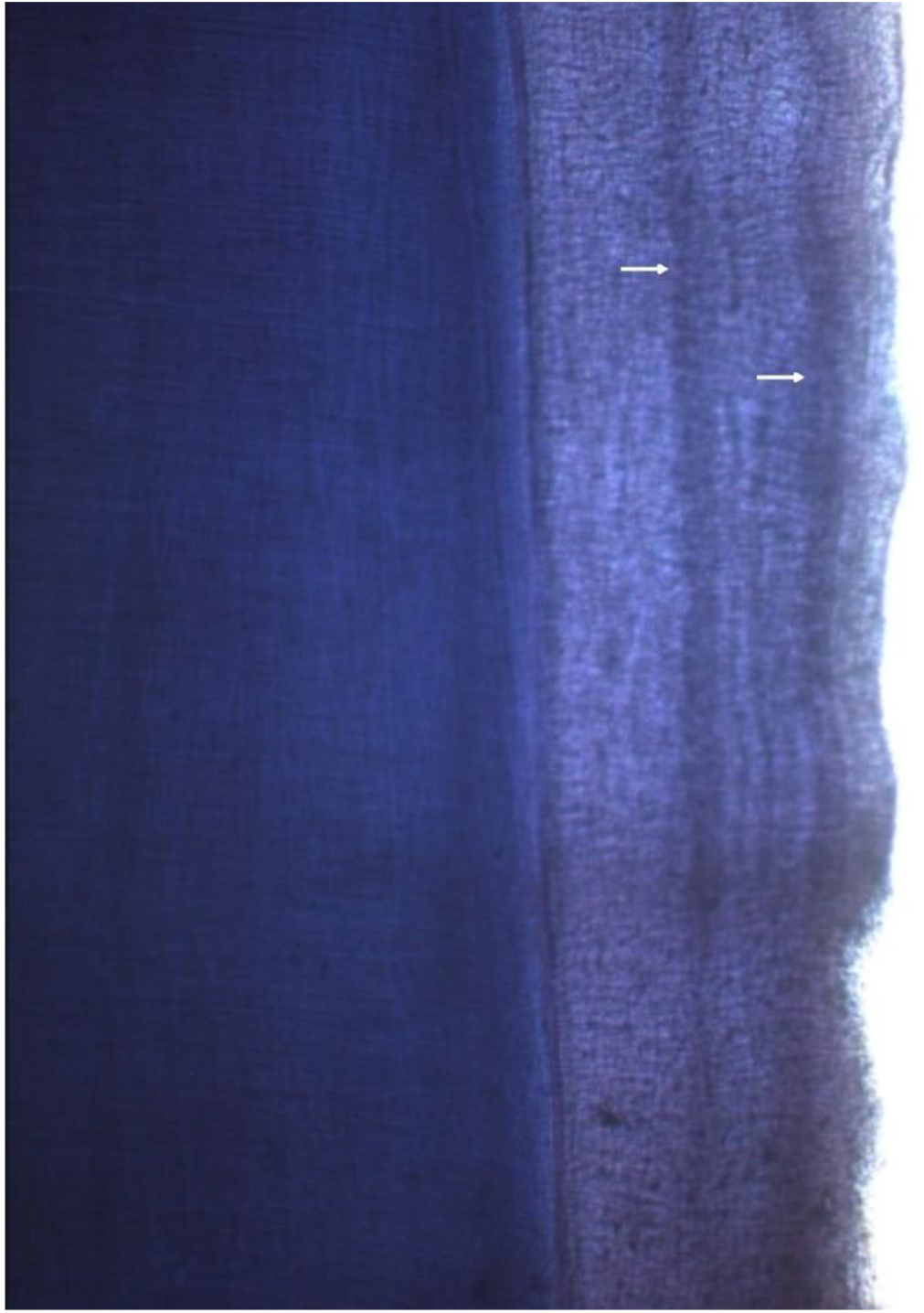
The arrows in the longitudinal section of I^3^ from seized tiger skeleton showing two cementum layers.

We recommend the validation of the current procedure while estimating age from cementum layer count. Matson et al. (1993) suggested two main tests for the validation cementum analysis technique for the estimation of age. These are the “blind” duplicate test when we use two or more teeth from an animal, and the second is using a tooth of known age for aging but without having the prior knowledge of its age. The teeth of known age were not available for both species. However, all incisor teeth in respective species showed clear and distinct common cementum layers. The periodontal membrane in all studied teeth confirms that all cementum layers were present in the longitudinal sections. Hence, we can say that the current method can show all cementum layers clearly and distinctly in incisor teeth. It may also make researchers more independent from relying on costly commercial services providing facilities worldwide. This technique can also be applied to other carnivore species.

## Acknowledgments

We gratefully acknowledge the support of the Director and Dean, Wildlife Institute of India (WII), Dehradun. This study was funded by a grant-in-aid of the Ministry of Environment, Forest and Climate Change, Government of India.

## References

Angerbjorn, A., Hersteinsson, P., Tannerfeldt, M. (2004). Arctic foxes: Consequences of resource predictability in the arctic fox—two life history strategies. In: MacDonald DW, Sillero-Zubiri C, eds. Biology and conservation of wild canids. New York: Oxford Univ Press. 2004: pp. 164–72.

Azorit, C., Analla, M., Hervás. J. et al. (2004). Aging through growth marks in teeth of Spanish red deer (*Cervus elaphus hispanicus*). Wildlife Soc. Bull. 32: 702–710.

Barthold, J.A., Loveridge, A.J., Macdonald, D.W., Packer, C., Colchero, F. (2016). Bayesian estimates of male and female African lion mortality for future use in population management. J Appl Ecol. 2016; 53:295–304. doi: 10.1111/1365-2664.12594.

Berube CH, Festa-Bianchet M, Jorgenson JT (1999). Individual differences, longevity, and reproductive senescence in bighorn ewes. Ecology 80:2555–2565

Binder, W. J. and Van Valkenburgh, B. (2010). A comparison of tooth wear and breakage in Rancho La Brea sabertooth cats and dire wolves across time. J. Verte. Paleon. 30: 255–261.

Cheater, A. (2006). Use of the upper second premolar for age determination of the African lion (*Panthera leo*) in sub-saharan Africa, for purposes of remote monitoring. – MSc thesis, Tshwane Univ. of Technology, Tshwane, South Africa.

Conover, M. (2002). Resolving human-wildlife conflicts: the science of wildlife damage management. Boca Raton, Florida: Lewis Publishers.

Craighead, J. J., Craighead F. C. & McCutchen, H. E. (1970). Age determination of grizzly bears from fourth premolar tooth sections. J. Wildlife Manage. 34: 353–363.

Creel, S., Mills, M.G.L., McNutt, J. W. (2004). African wild dogs: Demography and population dynamics of African wild dogs in three critical populations. In: Macdonald DW, Sillero-Zubiri C, eds. Biology and conservation of wild canids. New York: Oxford Univ Press. 2004: pp. 337–50.

Currier, M. J. P. (1983). Felis concolor. Mammalian Species.(200:): 1–7.

Douhard M, Loe LE, Stien A, Bonenfant C, Irvine RJ, Veiberg V, Ropstad E, Albon S (2016). The influence of weather conditions during gestation on life histories in a wild Arctic ungulate. Proc R Soc Biol Sci Ser B 283:20161760. https://doi.org/10.1098/rspb.2016.1760

Ferreira, S. & Funston, P. J. (2010). Age assignment to individual African lions. S. Afr. J. Wildl. Res. 40: 1–9.

Foresman, K. R. (2012). Carnivores in hand. In: Boitani L, Powell RA. 2012: eds. Carnivore ecology and conservation: A handbook of techniques. Techniques in ecology and conservation. New York: Oxford Univ Press. 2012: pp. 130–51.

Frank, L.G., Woodroffe, R., Ogada, M.O. (2005). People and predators in Laikipia District, Kenya. In: Woodroffe R, Thirgood S, Rabinowitz A, eds. People and Wildlife: conflict or coexistence? New York: Cambridge Univ Press. 2005. pp. 286–304.

Gipson, P. S. et al. (2000). Accuracy and precision of estimating age of gray wolves by tooth wear. J. Wildl. Manage. 64: 752–758.

Grue, H. & Jensen, B. (1979). Review of the formation of incremental lines in tooth cementum of terrestrial mammals. Dan. Rev. Game Biol. 11: 1–48

Hamlin K., Pac D.F. & Sime C.A. et al. (2000). Evaluating the accuracy of ages obtained by two methods for Montana ungulates. J. Wildlife Manage. 64: 441–449.

Harris, S. (1978). Age determination in the red fox (*Vulpes vulpes*) and evaluation of technique efficiency as applied to a sample of suburban foxes. J. Zool. Lond. 18: 91–117.

Hewison AJM, Gaillard JM (2001). Phenotypic quality and senescence affect different components of reproductive output in roe deer. J Anim Ecol 70:600–608

Johnston, D.H., Joachim, D.G., Bachmann, P. et al. (1987). Aging furbearers using tooth structure and biomarkers. In: Novak M., Baker

Klevezal, G. A. and Kleinenberg, S. E. (1967). Age determination of mammals by layered structure in teeth and bones. – Fish. Res. Board Can. Transl. Ser. no. 1024, Arct. Biol. Stn, QC.

Marks, S. A. & Erickson, P.W. (1966). Age determination in black bear. J. Wildlife Manage. 30: 389–410.

Matson, G., Daele, L.V., Goodwin, E. et al. (1993). A laboratory mannual for cementum agedetermination of Alaskan brown bear first premolar teeth. Matson’s Laboratory, Milltown, Montana.

Matson, G. M. (1981). Workbook for cementum analysis. – Matson’s Laboratory, Milltown, MT <www.matsonslab.com>.

Mazak. (1979). Der Tiger Panthera tigris. 2nd ed., Neue Brehm Bücherei 356. A. Ziemsen Verlag. Wittenberg Lutherstadt (GDR), 228 pp.

Mazák, Vratislav. (1981). Panthera tigris. MAMMALIAN SPECIES NO. 152, pp. 1–8, 3 figs. The American Society of Mammalogists.

Mbizah, M.M., Steenkamp, G. & Groom, R.J. (2016). Evaluation of the applicability of different age determination methods for estimating age of the endangered African wild dog (*Lycaon Pictus*). PLOS ONE 11: e0164676.

Miles, A. E. W. & Grigson, Caroline. (2003). *Colyer’s* Variations and diseases of the teeth of animals. Cambridge University Press.

Mundy, K.R.D. & Fuller, W.A. (1964). Age determination in grizzly bear. J. Wildlife Manage. 28: 863–866.

Nussey DH, Kruuk LEB, Donald A, Fowlie M, Clutton-Brock TH (2006). The rate of senescence in maternal performance increases with early-life fecundity in red deer. Ecol Lett 9:1342–1350.

Pelletier F, Moyes K, Clutton-Brock TH, Coulson T (2012). Decomposing variation in population growth into contributions from environment and phenotypes in an age-structured population. Proc R Soc Biol Sci Ser B 279:394–401. https://doi.org/10.1098/rspb.2011.0827

Schaller, G. B. (1972). The Serengeti lion: a study of predator–prey relations. Univ. of Chicago Press.

Schneider, K.M. (1959). Zum Zahndurchbruch des Lowen (*Panthera led*) nebst Bemerkungen iiber das Zahnen einiger andere Grosskatzen und der Hauskatze (*Felis catus*). Der Zoologische Garten, 22, 240–361. 339, 340.

Skalski J, Ryding K, Millspaugh J. 2005: Wildlife demography: Analysis of sex, age, and count data. Burlington, MA: Elsevier Inc.

Slaughter, B. H. et al. (1974). Eruption of cheek teeth in Insectivora and Carnivora. J. Mammal. 55: 115–125.

Smuts, G. L. et al. (1978). Age determination of the African lion (*Panthera leo*). Zool. Lond. 185: 115–146.

Spinage, C. A. (1976). Incremental cementum lines in the teeth of tropical African mammals. J. Zool., Lond. 178, I17–I3I.

Stander, P. E. (1997). Field age determination of leopards by tooth wear. Afr. J. Ecol. 35:156–161.

Veiberg, V., Nilsen, E.B., Rolandsen, C.M. et al. (2020). The accuracy and precision of age determination by dental cementum annuli in four northern cervids. Eur J Wildl Res 66, 91 (2020). https://doi.org/10.1007/s10344-020-01431-9

Vipin, Sharma V., Gupta S. K., Sharma, C. P., Sankar, K., Goyal, S. P. (2018). Development of a fast and low-cost age determination method in spotted deer, Axis axis. Folia Zool. – 67 (3-4): 186–197 DOI: 10.25225/fozo.v67.i3-4.a9.2018.

Weladji RB, Mysterud A, Holand Ø, Lenvik D (2002). Age-related reproductive effort in reindeer (Rangifer tarandus): evidence of senescence. Oecologia (Berl) 131:79–82

White, P. A., Ikanda, D., Ferrante, L., Chardonnet, P., Mesochina, P., Cameriere, R. (2016). Age Estimation of African Lions Panthera leo by Ratio of Tooth Areas. PLoS ONE 11(4): e0153648. doi:10.1371/journal.pone.0153648.

White, Paula, A., and Belant, Jerrold, L. (2016). Individual variation in dental characteristics for estimating age of African lions. Wildlife Biology 22: 71–77, 2016. doi:10.2981/wlb.00180

White, Paula A., and Belant, Jerrold L. 2016: Individual variation in dental characteristics for estimating age of African lions. Wildlife Biology 22: 71–77, 2016. doi: 10.2981/wlb.00180

Whitman, K. et al. (2004). Sustainable trophy hunting of African lions. Nature 428: 175–178.

Whitman, K. L. and Packer, C. (2007). A hunter’s guide to aging lions in eastern and southern Africa. Safari Press.

Willey, C.H. (1974). Aging black bears from first premolar tooth sections. J. Wildlife Manage. 38: 97–100

Williams, V.L., Newton, D.J., Loveridge, A.J., Macdonald, D.W. (2015). Bones of contention: An assessment of the South African trade in African lion Panthera leo bones and other body parts. Cambridge, UK and Wild CRU, Oxford, UK: TRAFFIC. 2015.

Zapata, S.C., Travaini, A. & Delibes, M. (1995). Comparacion entre varias tecnicas de estimacion de la edad en zorros, *Vulpes vulpes*, de Doñana (sur de la peninsula iberica). Doñana Acta Vertebrata 22: 29–50.

Zapata, S. C. et al. (1997). Age determination of Iberian lynx (*Lynx pardinus*) using canine radiograph and cementum annuli enumeration. Z. Saugetierkunde 62: 119–123.

